# An optimized pipeline for high-throughput bulk RNA-Seq deconvolution illustrates the impact of obesity and weight loss on cell composition of human adipose tissue

**DOI:** 10.1101/2024.09.23.614489

**Authors:** Cheehoon Ahn, Adeline Divoux, Mingqi Zhou, Marcus M Seldin, Lauren M Sparks, Katie L Whytock

## Abstract

Cellular heterogeneity of human adipose tissue, is linked to the pathophysiology of obesity and may impact the response to energy restriction and changes in fat mass. Here, we provide an optimized pipeline to estimate cellular composition in human abdominal subcutaneous adipose tissue (ASAT) from publicly available bulk RNA-Seq using signature profiles from our previously published full-length single nuclei (sn)RNA-Seq of the same depot. Individuals with obesity had greater proportions of macrophages and lower proportions of adipocyte sub-populations and vascular cells compared with lean individuals. Two months of diet-induced weight loss (DIWL) increased the estimated proportions of macrophages; however, two years of DIWL reduced the estimated proportions of macrophages, thereby suggesting a bi-phasic nature of cellular remodeling of ASAT during weight loss. Our optimized high-throughput pipeline facilitates the assessment of composition changes of highly characterized cell types in large numbers of ASAT samples using low-cost bulk RNA-Seq. Our data reveal novel changes in cellular heterogeneity and its association with cardiometabolic health in humans with obesity and following weight loss.

**Lead contact:** Katie Whytock

## INTRODUCTION

Adipose tissue is an important lipid storage and endocrine organ (1–3) that is highly heterogeneous where non-adipocytes compose more than 50% of the total cell population and reside within the stromal vascular fraction (e.g. stem cells, pre-adipocytes, vascular cells, and immune cells) (4, 5). While excess adiposity is associated with cardiometabolic disease progression (6, 7), increasing evidence suggests that altered cellular composition of adipose tissue – and not the sheer mass *per se* – is also tightly linked to, or even at the root of, cardiometabolic health complications that are often observed with obesity (8–10). Conversely, improvements in cardiometabolic health induced by weight loss are often accompanied by changes in the cellular composition of adipose tissue, particularly macrophages (11, 12). However, a comprehensive analysis of cell types that may be altered by weight loss is still lacking. Therefore, robust and accurate quantification of cell proportions in adipose tissue is paramount for understanding the etiology of cardiometabolic disease and optimizing its treatment.

Since the advent of transcriptomics, researchers have sought to deconvolute bulk transcriptomics to estimate cell type proportions. This approach has surged with the development of single cell (sc) and single nuclei (sn) RNA-Seq platforms that can more accurately quantify cellular composition and provide cell-specific transcriptomes to aid in bulk deconvolution. While sc/sn RNA-Seq remains the most accurate way to quantify cell composition in adipose tissue with minimal bias, the pipeline remains expensive and requires technical bench-work that restricts broad application. Recently, we published a full-length snRNA-Seq atlas of abdominal subcutaneous adipose tissue (ASAT) from a prospective cohort of older and younger adults balanced for sex and body mass index (BMI) (13). The full-length snRNA-Seq methodology provided the highest gene detection per nuclei in human adipose tissue to date, therefore providing an exemplary dataset for accurate bulk RNA-Seq deconvolution. While bulk RNA-Seq deconvolution using sc/snRNA-Seq profiles has previously been performed in human ASAT (9, 14, 15), to date no one has systematically identified which algorithm and signature matrix yields the most accurate results.

Here, we leverage our full-length snRNA-Seq human ASAT dataset to optimize a pipeline to deconvolute bulk RNA-Seq datasets and determine how cellular heterogeneity of ASAT may be altered with both obesity and weight loss. Understanding the impact of one of the most cost-effective and commonly prescribed interventions to treat obesity-related cardiometabolic health complications may lead to advanced weight loss strategies and cellular targets for improving cardiometabolic health outcomes.

## RESULTS

### Assessment of deconvolution algorithms

Given there are currently several popular deconvolution algorithms, we aimed to assess which algorithm had the ability to 1) detect every cell type that we previously reported (13), from a large bulk RNA-Seq data set and 2) how accurately we could deconvolute a pseudobulk RNA-Seq data set, which is a bulk-like profile where gene expression data from individual nuclei are aggregated for each sample with known cell type proportions. In our preliminary testing, we compared popular algorithms for their capability of detecting each cell type in the majority of samples from a large bulk RNA-Seq data set (METabolic Syndrome In Men; METSIM cohort (16) **Figure 1**). We also reasoned that for the algorithm to be effective it should be able to detect at least 25% of adipocytes in the majority of these samples (17). The results show that certain algorithms, despite the different gene composition iterations, were never reliable in detecting certain cell types. For example, when using all detected genes (‘All Genes’) and top 5000 highly variable genes (HVG) – which were used to cluster cell types in the original snRNAseq data (13) – rls and qprogwc always underestimated vascular cells. nnls (operated through ADAPTS or granulator) did not identify adipocytes in a large portion of samples (**Figure 1A, S1A**). MuSiC-weighted consistently underpredicted stem and vascular proportions whereas MuSiC – all gene underpredicted macrophages and pre-adipocyte proportions (**Figure 1A, S1A)**. While EPIC was able to detect all cell types, the proportion of these certain cell types in samples was extremely low (<1%) and therefore was not a completely reliable detection. Some algorithms such as ols, qprog, DCQ, proportionsInAdmixture, and DeconRNASeq were able to detect adipocytes in a large proportion of samples. However, these adipocyte estimates had an extremely low proportion of samples that had >25% of adipocyte estimated (**Figure 1B**). It was notable that dtangle was able to detect all cell type in every sample when all genes or 5000HVG signature was used (**Figure 1A, S1A)**. Furthermore, dtangle consistently detected at least 25% of adipocytes in nearly 100% of samples (**Figure 1B, S1B**).

**Figure 1.**
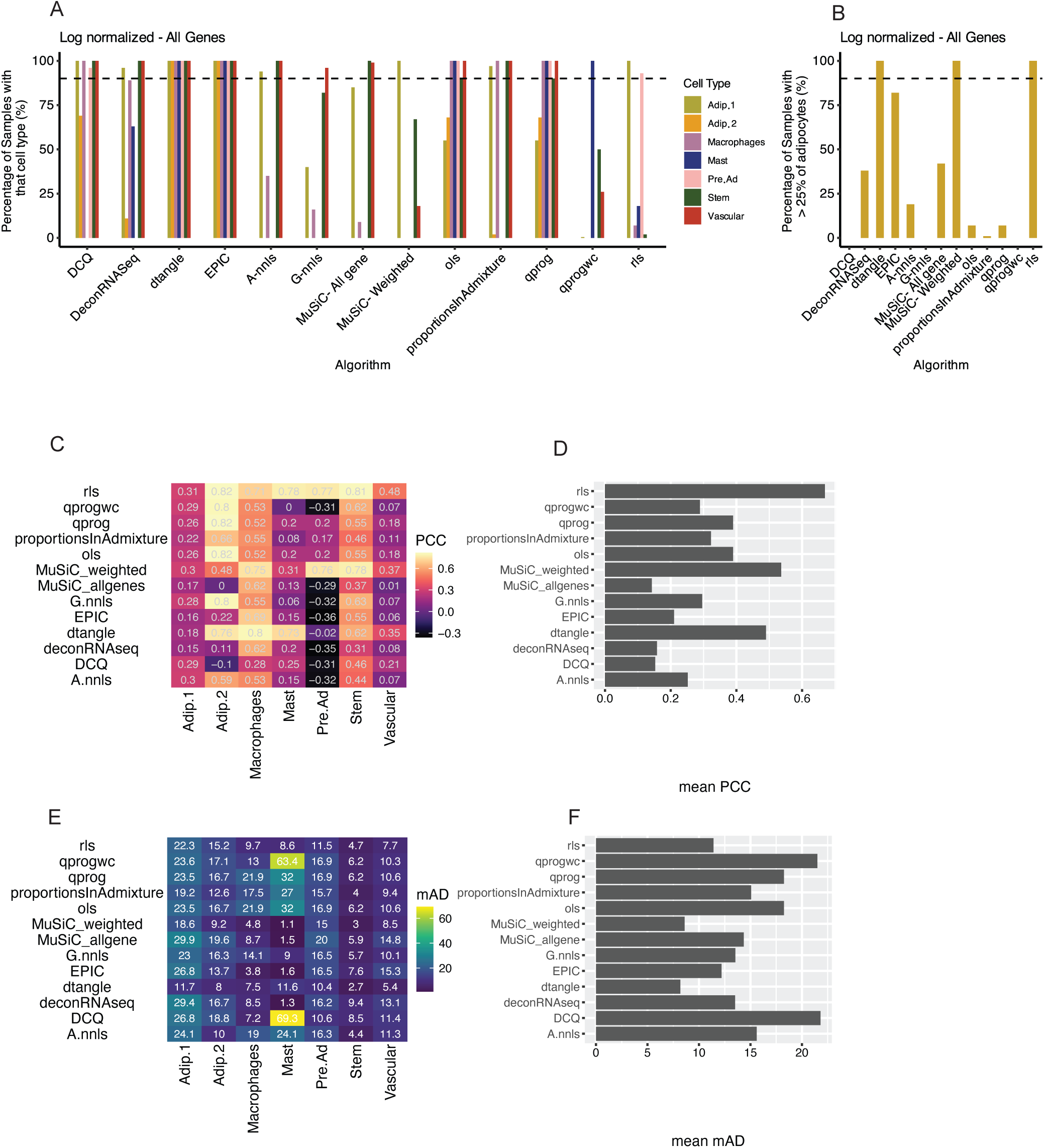
Assessment of deconvolution algorithms. (A) The percentage of samples from the METSIM cohort that are estimated to contain that cell type when predicted with different deconvolution algorithms using log normalized signature matrix from all detected genes. (B) The percentage of samples from the METSIM cohort that are estimated to have >25% of adipocytes predicted with different deconvolution algorithms using log normalized signature matrix from all detected genes. (C) The pearsons correlation coefficient (PCC) comparing estimated cell proportions from pseudobulk RNA-Seq data against quantified cell proportions from snRNA-Seq for different algorithms for each cell type (D) and the average PCC score across all cell types. (E) The mean absolute deviation (mAD) between estimated cell proportions pseudobulk RNA-Seq data and quantified cell proportions from snRNA-Seq for different algorithms for each cell type (F) and the average mAD score across all cell types. If box is grey the algorithm did not estimate that cell type and PCC therefore cannot be quantified. HVG, Highly variable genes.

In order to assess the effectiveness of different deconvolution algorithms, we compared estimated cell proportions from a pseudobulk data-set against actual cell-type proportions quantified with snRNA-Seq. Pearsons’s correlation coefficient (−1 to 1) and mean absolute deviance (mAD) (0–100) were used to assess how accurately the deconvolution estimated cell type proportions, with a PCC of 1 and mAD of 0 indicating a completely accurate prediction of a given cell type (**Figure 1C-F).** Overall dtangle, rls and MuSiC-weighted had the highest PCC values (**Figure 1C-D**). dtangle and MuSiC-weighted had the lowest average mAD out of all the algorithms that were able to detect every cell type (**Figure 1E-F**).

### Signature Matrix Optimization

Due to dtangle being the best-performing algorithm from the bulk and pseudobulk RNA-Seq deconvolution assessments, we further sought to optimize this specific algorithm.

In adipose tissue biology, macrophage content increases proportionally to increases in adipose tissue mass (18). We therefore reasoned that when deconvoluting bulk RNA-Seq data, we should see a positive correlation between macrophage proportion and adiposity (i.e., BMI and waist-to-hip ratio (WHR)). We ran different iterations of gene signatures to assess which gave us the best correlation between macrophage proportion and BMI or WHR. We tested; all genes detected, and different iterations of HVG (2000, 3000, 4000, 5000, 6000, 7000, 8000, 9000, 10000). Every gene signature was able to detect every cell type in every sample (**Figure 2A**). However, ‘All genes’ *signature matrix* predicted similar proportions of macrophages across all samples and therefore there was no correlation between percentage of macrophage proportion and WHR (**Figure 2B, D**) or BMI (**Figure S2A**). This highlights the need to optimize a gene signature when performing deconvolution rather than using all of the genes available. The 6000 HVG had the highest correlation between both WHR and BMI with estimated proportion of macrophages, although 5000 HVG, 4000 HVG, and 3000 HVG elicited similar results (**Figure 2B, C, S2A**).

**Figure 2.**
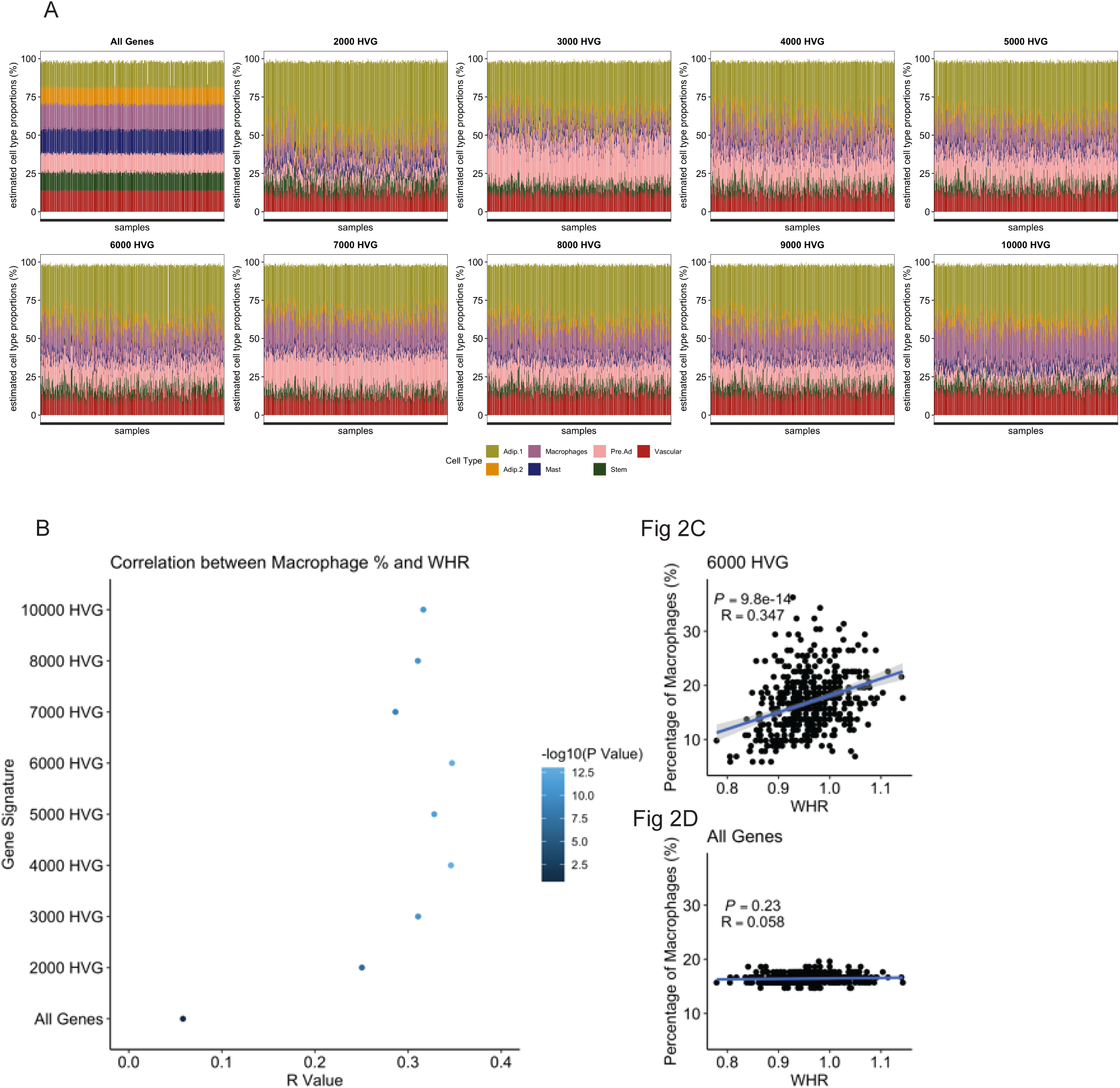
Correlation between estimated macrophage proportions and BMI or WHR using different gene signatures to subset the signature matrix. (A) Estimated cell type proportions of each METSIM RNA-Seq samples using different HVG to modify the signature matrix. Algorithm dtangle was used and the data was log normalized. (B) R value and –log10(p value) for estimated macrophage proportion and WHR for the METSIM RNA-Seq data. (C) Scatterplot showing the estimated macrophage proportion and WHR for the 6000 HVG or (D) all genes signature.

We further explored whether the snRNAseq-derived gene signature matrix and 6000 HVG list were influenced by the age of the participants as our snRNAseq reference data comprises samples from 10 older (≥ 65 years)) and 10 younger (≤ 30 years) participants (13). By applying age-group specific gene signature matrices and age-group specific 6000 HVG lists to deconvolute METSIM bulk RNA-seq data, we found that macrophages were overestimated while pre-adipocytes were underestimated when using gene signature matrix and 6000 HVG list from younger adults compared with when using those from older adults (**Figure S2B-C**). In addition to our main signature matrix and 6000 HVG from the age-group-integrated data, we provide age-group-specific data (**Supporting information**). Using the 6000 HVG from the two different age groups, we found that 4762 genes among 6000 HVG (65.9%) were shared by both groups, suggesting that these 4762 genes may represent ‘age-neutral’ genes (**Figure S2D**). There was a remarkably tight correlation between the estimated proportions of each cell type from METSIM bulk RNA-seq data using the age-group-integrated 6000 HVG and ‘age-neutral’ 4762 HVG (0.94 < R < 0.99) (**Figure S2E**), indicating that the initial 6000 HVG we tested may robustly estimate ASAT cell type proportions with minimal bias by age.

### Deconvolution reveals distinct ASAT heterogeneity in obesity

To further examine how ASAT cellular heterogeneity may be implicated in obesity and cardiometabolic health, we deconvoluted ASAT bulk RNA-seq data from previously published reports that collected ASAT from cohorts of lean and individuals with obesity that represented a wide range of age; adolescents (<18 years; obesity defined as 97^th^ percentile BMI; “LCAT cohort”) (**Figure 3A**) (19), young and middle-aged adults (18-55 years; “Petersen et al., 2024”) (**Figure 3B**) (20), and older adults (55-70years; “MD lipolysis cohorts”) (**Figure 3C**) (21). In addition to cohorts of adults with obesity who had relatively healthier cardiometabolic health traits (i.e. Metabolically Healthy Obese, MHO and Obese with insulin resistance, Obese-IR), two studies had another cohort that had impaired cardiometabolic health traits. For example, Metabolically Unhealthy Obese (MUO) had prediabetes, hepatic steatosis, and whole-body insulin resistance (**Figure 3B**) and Obese with Type 2 diabetes (Obese-T2D) were recently diagnosed with T2D by the time they were recruited (**Figure 3C**). Importantly, these cohorts were matched to their healthier cohorts with obesity by sex, age, and adiposity (see more details in ‘*Human studies and deconvolution analysis’*). Consistently observed across different age groups, our deconvolution analysis estimated a higher proportion of macrophages and a lower proportion of vascular cells in individuals with obesity compared with lean cohorts, aligning with the adipose tissue abnormalities (i.e., macrophage infiltration and capillary rarefaction) commonly observed in obesity (18, 22, 23) (**Figures 3A-C**). We previously characterized two adipocyte populations using snRNA-Seq (13). Adipocyte 1 was characterized by an upregulation of genes related to anti-oxidation (*GPX1 & GPX4*) and pathways related to complement, oxidative phosphorylation and Srp dependent translational protein targeting to membrane and was labeled as ‘anti-oxidative’, while adipocyte 2 was labeled as ‘insulin-responsive’ adipocyte, demonstrated by upregulation of genes related to suppression of lipolysis (*PDE3B*), lipid metabolism (*ABCA5),* and of pathways related to insulin receptor signaling cascade (13). Interestingly, the estimated proportions of both adipocytes were also lower in individuals with obesity compared with lean individuals (**Figures 3A-C**). The estimated proportions of pre-adipocytes, stem cells, and mast cells were comparable between the individuals with obesity compared with lean (**Figures 3A-C**).

**Figure 3.**
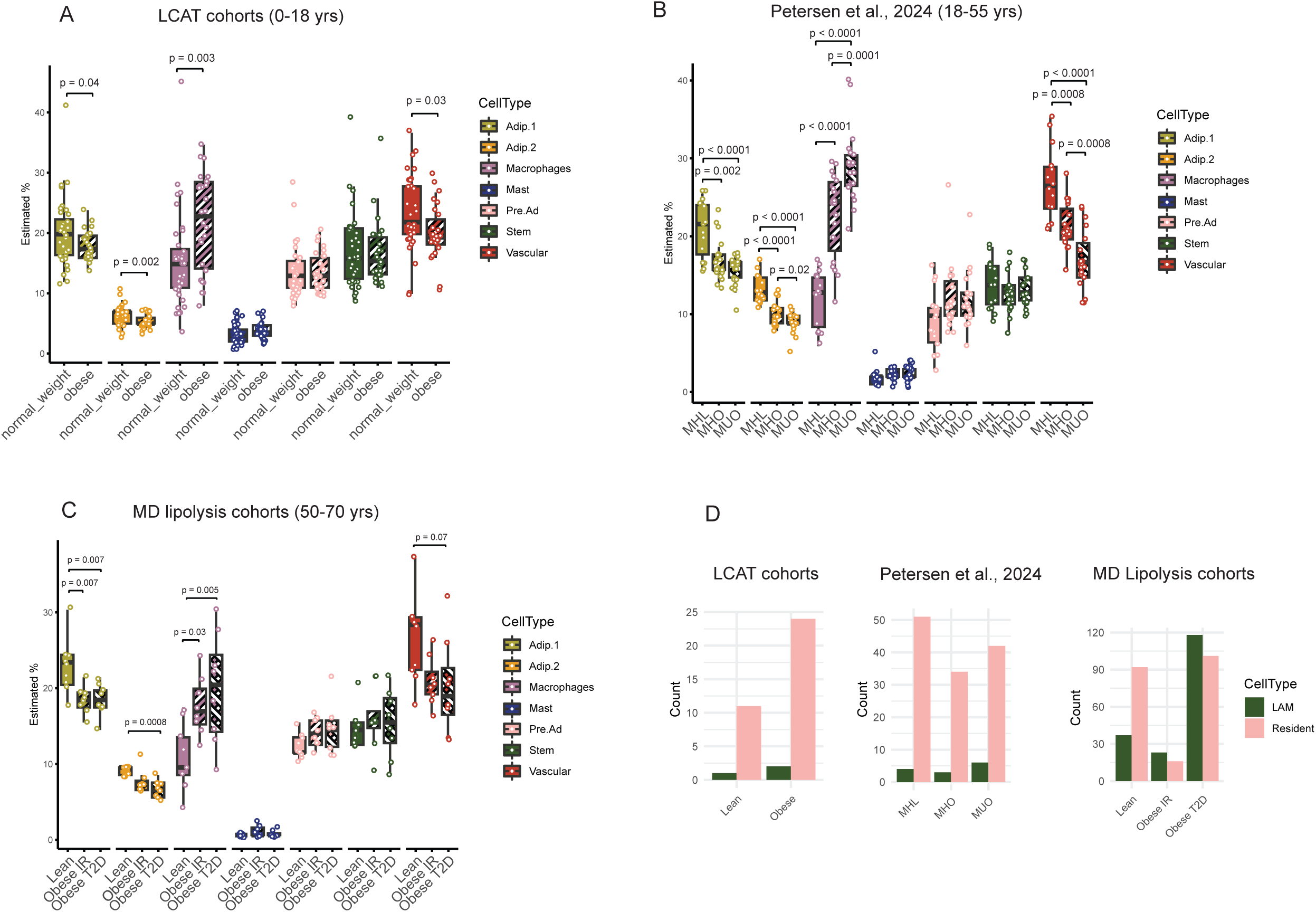
Estimated ASAT cell type proportions from cross-sectional obesity studies. (A) Deconvoluted ASAT cell type proportions from lean and obese cohorts in LCAT study (19). Sample size: Normal weight = 35, Obese = 26. (B) Deconvoluted ASAT cell type proportions from cohorts of MHL, MHO, and MUO (20). Post-hoc Tukey HSD was used for cell types with significant ANOVA group differences. Sample size: MHL= 15, MHO = 19, MUO = 19. (C) Deconvoluted ASAT cell type proportions from cohorts of lean, Obese with IR, and Obese with T2D in MD lipolysis cohorts (21). Tukey HSD was used for post hoc analysis of cell types with significant ANOVA group differences. Sample size: Lean = 9, Obese IR = 8, Obese T2D = 8. (D) Number of marker genes for LAM or resident macrophage that are significantly associated with the estimate proportion of macrophages in each study (19–21). LCAT; Leipzig Childhood Adipose Tissue, MHL; Metabolically Healthy Lean, MHO: Metabolically Healthy Obese, MUO; Metabolically Unhealthy Obese, HSD; Honestly Significant Difference, IR; Insulin resistance, T2D; Type 2 Diabetes. LAM; Lipid associated macrophage.

### Deconvolution reveals ASAT cell types that are linked with poor cardiometabolic health

Macrophage infiltration and capillary rarefaction in ASAT are often associated with unfavorable cardiometabolic health in obesity, but the interpretation of their direct link can be confounded by increasing ASAT mass (7, 24). We investigated whether the estimated cellular composition differs between adults with obesity who have cardiometabolic health complications (i.e., MUO and Obese-T2D) and those well-matched individuals with obesity who are relatively healthy (i.e., MHO and Obese-IR). Compared with the MHO group, the MUO group was estimated to have a higher proportion of macrophages (p=0.0001) and a lower estimated proportion of adipocyte 2 and vascular cells (p=0.02 and p=0.0008, respectively) (**Figure 3B**). In the MD lipolysis cohorts, while the estimated proportion of adipocyte 2 was not different between older lean vs. older adults with Obese-IR, it was significantly lower in older adults with Obese T2D when compared with lean (p=0.0008) (**Figure 3C**). There was a trend for lower estimated proportion of vascular cells in Obese T2D compared to lean (p=0.07) (**Figure 3C**), collectively suggesting that alterations in macrophage and vascular cell populations may be directly linked with impairments in cardiometabolic health. Intriguingly, our findings further suggest that a lower proportion of insulin-responsive adipocytes may also be implicated in adverse cardiometabolic health outcomes.

We then used a ‘gene inference’ approach to understand whether or how specific phenotypes of adipose tissue macrophages (e.g., M1-like pro-inflammatory vs. M2-like anti-inflammatory) may be altered with obesity. By using marker genes for M1-like lipid-associated macrophage (LAM) (n=317 genes) or M2-like resident macrophages (n=2724 genes) that we previously acquired from our parent snRNA-Seq data (13), we conducted a correlation analysis between marker gene expressions and estimated proportions of macrophages. The number of resident macrophage marker genes that are significantly and positively correlated with the estimated proportion of macrophages was lower in adults with obesity (i.e., MHO and Obese IR) compared to lean (**Figure 3D**), and higher in adolescents with obesity compared with lean adolescents (**Figure 3D**). These data suggest a possibility that adipose tissue macrophage polarization may be differentially regulated with obesity in youth. Notably, the number of associated marker genes for LAM was higher in adults with obesity and impaired cardiometabolic health (i.e., MUO and Obese T2D) compared with their well-matched obese groups (**Figure 3D**), aligning with the authors’ finding of higher markers of whole-body and local inflammation in MUO and Obese T2D (20, 21).

### Diet-induced weight loss modifies ASAT cell type proportions

Diet-induced weight loss (DIWL) is known to improve cardiometabolic health, often accompanying alterations in the ASAT microenvironment that include reduced inflammation and improved lipid metabolism (25–28). However, it is unclear whether cell type composition in ASAT can be modified by DIWL and may underlie the improved health outcomes observed with DIWL. We deconvoluted ASAT bulk RNA-Seq data from several dietary intervention studies that induced weight loss of ∼10% (range: 8∼11%). Comprehensive Assessment of the Long-term Effects of Reducing Energy Intake (CALERIE) Study is a randomized clinical trial that examined the effects of 12 and 24 months of 25% caloric restriction (CR) in humans without obesity (29). Compared with the control group (AL, Ad Libitum), participants in the CR group lost ∼11% of weight (∼6kg fat mass) in response to 12 months of CR, which was maintained at 24 months (**Table S1**). Our deconvolution analysis suggested that the proportion of macrophages was significantly reduced after 12 and 24 months of CR compared to baseline (p=0.002 and p=0.003, respectively) (**Figure 4A**). In concordance with this finding, there was a trend of positive correlation between the change in fat mass (Δ kg) and the change in estimated macrophage proportion (Δ %) from baseline to month 12 (p=0.07) (**Figure 4B**) i.e. a decrease in fat mass (kg) correlates with decreases in estimated macrophage proportion. Interestingly, we did not observe a decrease in estimated macrophage proportion in the other two studies where short-term caloric restriction was prescribed to individuals with overweight/obesity. In the Diet, Obesity, and Genes (DiOGene) study, 220 adults with overweight/obesity (Age: 18-65 years, BMI: 27-45kg/m^2^) underwent an 8-week low-calorie diet (LCD), all achieving at least 8% weight loss by the time of their post-ASAT sample collection (**Table S2**) (26). In agreement with the findings of upregulated expression of macrophage genes in these participants (26), our deconvolution analysis suggested an increased proportion of macrophages in response to eight weeks of LCD (p=5.64e-13) (**Figure 4C**). Conversely, the estimated proportion of adipocyte 1, adipocyte 2, and vascular cells was reduced (p=2.24e-17, 8.32e-5, 0.016, respectively) with a slight increase in estimated stem cell proportion (p=0.014) (**Figure 4C**). Similarly, we observed a trend of increased estimated proportion of macrophages (p=0.062) and reduced estimated proportion of adipocyte 1 and adipocyte 2 (p=0.021 and 0.013, respectively) from a small cohort of women with obesity (n=10, Age: 61±4 years, BMI: 39±3kg/m^2^) who rapidly lost ∼10% of their weight through very low-calorie diet (VLCD) over 6.6±2.2 weeks (**Figure 4D**, **Table S2**) (27). Interestingly, the authors of this study reported an increased density of crown-like structures – which are aggregates of macrophages – in ASAT after VLCD, which aligns with our findings from the deconvolution analysis.

**Figure 4.**
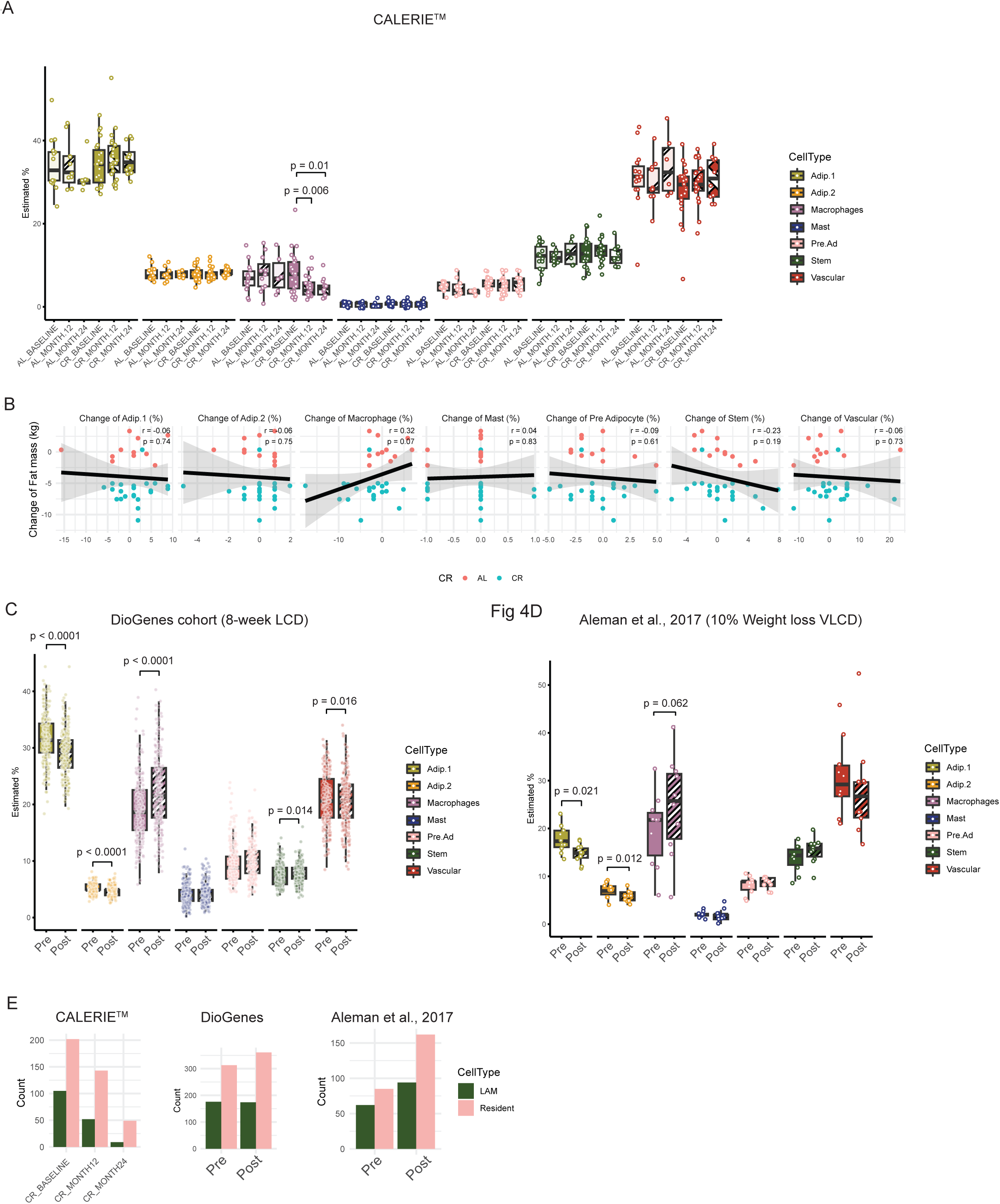
Estimated ASAT cell type proportions from longitudinal DIWL studies. (A) Deconvoluted ASAT cell type proportions from CALERIE^TM^ cohorts. Least square means was used for post hoc analysis of macrophage in the CR group. Sample size; AL-Baseline, n=14, AL-Month 12, n=11, AL-Month 24, n=6; CR-Baseline, n=23; CR-Month12, n=23; CR-Month 24, n=12. (B) Correlation between change of fat mass (Δ kg) and change of cell type proportions (Δ %) in CALERI^TM^ participants who completed 12-month intervention. (C) Deconvoluted ASAT cell type proportions from DioGenes cohorts who underwent 8-week low-calorie diet using meal replacement product (26). Sample size = 220. (D) Deconvoluted ASAT cell type proportions from women with obesity who achieved 10% weight loss by very low-calorie diet (27). Sample size = 10. (E) Number of marker genes for LAM or resident macrophage that are significantly associated with the estimate proportion of macrophages in each study (26, 27, 29). DIWL, Diet-induced weight loss; AL, Ad Libitum; CR, Calorie Restriction.

Additionally, the number of significantly associated marker genes for both LAM and resident macrophages with estimated proportions of macrophages was reduced by 12 and 24 months of CR in CALERIE (**Figure 4E**). However, the magnitude of reduction was greater in LAM marker genes compared with resident macrophage marker genes, leaving less than 10% of significantly associated genes after 2 years of CR (**Figure 4E**). Although we observed an increased estimated proportion of macrophages in DioGenes cohorts and Aleman et al., 2017 (**Figures 4C, D**), our ‘gene inference’ approach showed a greater increase in the number of resident macrophage marker genes that are associated with estimated macrophage proportion, compared with LAM marker genes. This finding suggests that the increased estimated proportion of macrophages by short-term DIWL in adults with obesity may have been driven by an increased abundance of resident macrophages, which has been reported to buffer lipids and counteract inflammatory responses, thereby contributing to appropriate and favorable adipose tissue remodeling (30).

## DISCUSSION

Using publicly available algorithms and a refined the reference data derived from our published full-length snRNA-Seq (i.e., gene signature matrix and 6000HVG (13)), we established a deconvolution pipeline that robustly estimates ASAT cell type proportions from bulk RNA-Seq data that requires less cost and labor compared with sc or snRNA-seq. Using this algorithm, we identify cellular heterogeneity in obesity, which is characterized by higher proportions of macrophages, and lower proportions of adipocytes and vascular cells. Additionally, altered abundance of some cell types, such as insulin-responsive adipocytes, macrophages, and vascular cells may directly underlie the impaired cardiometabolic health in individuals with obesity. We further show a dynamic change in cell type proportions in response to DIWL, which may play an important role in mediating some of the health benefits conferred by DIWL.

Increased adipose tissue macrophage infiltration and capillary rarefaction are hallmarks of excessive adiposity that are tightly associated with functional abnormalities in adipose tissue (8, 22, 23, 31). Our findings of higher estimated proportions of macrophages and lower estimated proportions of vascular cells in individuals with obesity confirm this notion and further suggest that these ASAT abnormalities are linked with cardiometabolic health complications independent of ASAT mass. Additionally, higher estimated proportions of macrophages in MUO and Obese-T2D were associated with more marker genes of LAMs compared with MHO and Obese-IR respectively, suggesting that alterations in both macrophage abundance and phenotype may be directly linked with cardiometabolic health complications in middle-aged and older adults. However, this phenotypical switch of macrophages towards a pro-inflammatory type during the progression of obesity may not apply in adolescent populations. We found that the estimated proportions of macrophages in adolescents with obesity in the LCAT cohorts were more associated with marker genes for anti-inflammatory type resident macrophages. We speculate that ASAT in adolescents with obesity may require a higher abundance of resident macrophages because it is under continuous tissue expansion and remodeling at such a young age (32–34).

Our deconvolution outcomes suggested a possibility of a lower proportion of mature adipocytes in obesity, which serve as primary storage for excess lipid. Lower proportions of adipocytes do not translate to lower number of total adipocytes when comparing individuals with obesity to lean individuals. For example, the total number of adipocytes was estimated to be more than 2.5-fold higher in adolescents with obesity compared to lean adolescents in the LCAT cohorts (35). Therefore, while total adipocytes in adipose tissue may be increased with increasing adiposity, its relative abundance to other cell types appears to be reduced. Our novel finding that the estimated proportion of insulin-responsive adipocytes (adipocyte 2), characterized by upregulated genes involved in lipid storage/metabolism and insulin signaling (13), was lower in individuals with obesity and impaired cardiometabolic health (i.e., MUO and Obese-T2D) compared with matched individuals with obesity who are relatively healthy (i.e., MHO and Obese-IR) indicates a tight relationship between adipocyte heterogeneity and cardiometabolic health. Although the potential mechanistic link bridging these two is unclear, the lower estimated proportion of adipocyte 2 in MUO was paralleled with significantly higher ectopic fat mass (i.e., intra-abdominal adipose tissue and intrahepatic triglyceride content) (20), which is indicative of impaired lipid storage capacity of the ASAT (36, 37). We therefore speculate that the lower abundance of adipocyte 2 may have contributed to the limited lipid storage capacity of ASAT in MUO, resulting in the accumulation of ectopic fat, which can cause tissue-specific and whole-body insulin resistance (38).

Although DIWL-mediated attenuation of systemic inflammation has been commonly reported in individuals with overweight/obesity (39, 40), the CALERIE study was the first to reveal that reduced markers of systemic inflammation after one to two years of CR in individuals without obesity (41). We found a reduced estimated proportion of ASAT macrophages after one and two years of CR in the CALERIE cohort which may explain the reduced systemic inflammation.

Reduced proportions of macrophages by one and two years of CR was also paralleled by a vast reduction of the number of associated marker genes for LAM, suggesting a phenotypical switch of macrophages in addition to the change of abundance. Conversely, the increased estimated proportion of ASAT macrophages in response to DIWL from DioGenes cohorts and participants in Aleman et al (27) indicates that cellular remodeling during DIWL may vary significantly depending on specific conditions and contexts. These findings support the ‘biphasic’ responses in adipose tissue inflammation during weight loss, where markers of macrophages and inflammation are increased during early weight loss then followed by a larger reduction as weight loss sustains (11, 25, 42). Perhaps, our view is that weight loss may improve lipolytic capacity of adipocytes that in turn induces macrophage recruitment (11), and our gene inference data suggest that this rise may be driven by an increased abundance of M2-like resident macrophages, indicating a potentially favorable adaptation (43).

Weight loss induces extensive morphological and functional remodeling of adipocytes, which includes reduced adipocyte size and restoration of lipid metabolism in obesity (11, 25, 44). Our finding of a reduced estimated proportion of adipocytes after short-term DIWL in adults with obesity (DioGenes and Aleman et al.) suggests a potential modification in adipocyte ‘abundance’ by weight loss. While this may simply be a reciprocal shift in proportion due to an increased proportion of other cell types (e.g., macrophage and stem cell), we cannot rule out the possibility that this may reflect changes in adipocyte turnover (i.e., net balance between adipocyte formation and deletion). Weight loss may inhibit adipogenesis (45) which may contribute to the negative balance of adipocyte abundance. However, many studies suggest the opposite (46, 47), and it is unlikely that the reduced rate of adipogenesis was translated into a meaningful reduction in adipocyte abundance in a relatively short period (∼8 weeks) given the slow rate of adipogenesis in humans (34). Alternatively, increased adipocyte apoptosis may have driven cellular turnover during weight loss (48). Interestingly, it was previously demonstrated that removal of adipocytes by apoptosis recruited M2-like macrophages into adipose tissue in mice (49), which may also explain the increased association of resident macrophage marker genes with increased estimated proportion of macrophages in response to short-term DIWL.

Although our study enabled robust estimation of cell type proportions in human ASAT from bulk RNA-Seq data, it is important to acknowledge some limitations that are inherent to *in silico* analysis. Since the bulk RNA-Seq datasets we used were derived from other authors, we could not directly validate our deconvolution analysis with the original tissue samples. However, many of our cell type estimations align with findings from original articles. For example, our findings of higher macrophage proportion in adolescents with obesity and after short-term VLCD in adults with obesity parallel with the higher macrophage immunostaining in LCAT cohorts with obesity and increased CLS after rapid VLCD in adults with obesity from Aleman et al. respectively, supporting the robustness and accuracy of our deconvolution pipeline. Additionally, while the full-length snRNA-Seq data from which we generated reference data has a superior gene coverage rate per nuclei compared with conventional single-end (i.e., 3’-end) snRNA-Seq techniques, subtypes of macrophages were not defined at the parent level, preventing direct estimation of those cell populations. However, by associating marker genes of LAM and resident macrophage acquired from a secondary cell clustering analysis (13) with estimated proportion of macrophages, we show that macrophage phenotypes may be altered by obesity and DIWL. Furthermore, the intriguing findings that the abundance of certain cell types (i.e., adipocyte 2, macrophage, and vascular cell) may be associated with the progression of cardiometabolic health complications, yet the direct linkage remains inconclusive.

In summary, our findings indicate that compared with lean individuals, those with obesity exhibit distinct cellular heterogeneity in ASAT, and further alterations in cell type proportions are tightly linked with impaired cardiometabolic health. Moreover, DIWL can induce dynamic alterations in ASAT cell type proportions, potentially contributing to the improved cardiometabolic health.

Overall, our work expands the understanding of adipose tissue cellular heterogeneity implicated in cardiometabolic health and weight loss interventions by providing an optimized computational deconvolution pipeline that can be easily used to estimate cell type proportions in human ASAT.

## METHODS

### BIOINFORMATIC ANALYSES

#### DATA GENERATION

Source data was generated in Seurat V4.4.0 with SeuratObject V4.1.4 using a previously published data set (13). Seurat objects from each individual sample were merged into 1 large seurat object before being split into a list of seurat objects based on individual samples. SCTtransform with glmGamPoi was performed on this seurat list. Highly Variable Genes (HVG) were determined by performing SelectIntegrationFeatures() on the SCTransformed seurat list.

A signature matrix was generated using AggregateExpression() and aggregating SCTtransformed counts for each cell type across all participants. The signature matrix was then filtered by HVG list and used for subsequent deconvolution analyses. A pseudobulk RNA-Seq data set was generated by using AggregateExpression() and aggregating SCTtransformed counts for each sample across all cell types.

#### PIPELINE OPTIMIZATION

We initially compared popular deconvolution algorithms that were either operated through R package granulator (50) (ols-ordinary least squares, qprog – quadratic programming without constraints, rls-re-weighted least squares (51), qprogwc-quadratic programming with non-negativity and sum-to-one constraint (52), nnls-non negative least squares (53) and dtangle (54)) or R package ADAPTS (55) (DCQ-Digitial cell quantifier (56), deconRNASeq (52), nnls-non negative least squares and ProportionsInAMixutre) and from stand alone R Packages MuSiC (weighted and All Genes) (57) and EPIC (58). As nnls algorithm was included in both granulator and ADAPTS we used both and assigned an A for Adapts and G for Granulator to identify which package it was performed on.

##### Algorithms comparison

We tested each algorithms capability of detecting each cell type and detecting at least 25% of adipocytes in the majority of samples from a large bulk RNA-Seq data set (METabolic Syndrome In Men; METSIM cohort, GSE135134 (16)). To ensure we were not prematurely dismissive of a certain algorithm we tested the deconvolution using; the full gene signature **(Figure S1A, B),** and using a signature matrix that used 5000 highly variable genes (named 5000 HVG) that was used for clustering in the original snRNA-Seq analyses. We reasoned that these 5000 HVG dictated the clustering of cells and therefore would be an optimal initial signature to detect cell types in bulk RNA-Seq data. For deconvolution using algorithms from granulator, function deconvolute() was performed with the TPM-normalized bulk RNA-Seq data and the SCTransform signature matrices. For deconvolution using algorithms using ADAPTS, functions estCellPercent.nnls(), estCellPercent.DCQ(), est.CellPercent.proportionsInAMixture(), estCellPercentDeconRNASeq() were performed with the TPM-normalized bulk RNA-Seq data and the SCTransform signature matrices. For deconvolution using MuSiC the integrated Seurat Object was first converted to a SingleCellExperiment with function as.SingleCellExperiment(). The TPM-normalized bulk RNA-Seq data was converted to an expression matrix with ExpressionSet() and exprs() and then music_prop() was performed with either all genes or with markers argument set to the 5000 HVG. For deconvolution using EPIC function EPIC() was performed on the TPM normalized bulk RNA-Seq data and the SCTransform signature matrices. For each cell type the percentage of samples estimated to have that cell type was calculated, in addition to the percentage of samples estimated to have at least 25% of adipocytes (adipocyte_1 and adipocyte_2 combined).

##### Pseudobulk assessment

In order to assess the effectiveness of different deconvolution algorithms we compared estimated cell proportions from a pseudobulk data-set against actual cell-type proportions quantified with snRNA-Seq. In this instance deconvolution was performed with the above functions and with the 5000HVG signature matrix but instead with a pseudobulk RNA-Seq data that was generated from aggregated SCTtransformed counts for each sample across all cell types. Principle correlation coefficient (−1 to 1) and mean absolute deviance (mAD) (0–100) were used to assess how accurately the deconvolution estimated cell type proportions. MAD was calculated as the sum of the absolute differences between the predicted proportion from the actual proportion divided by the by the total number of samples (n = 20).

##### Signature Matrix Optimization

Using the SelectIntegrationFeatures() we generated iterations of HVG lists (2000, 3000, 4000, 5000, 6000, 7000, 8000, 9000, 10000) and filtered the signature matrix to these gene lists creating a list of signatures matrices. Deconvolution was then performed using deconvolute() with just dtangle algorithm, using the list of signature matrices and the TPM normalized bulk RNA-Seq METSim data. Estimated Macrophage proportions were correlated to BMI and WHR (59), using function bicorAndPvalue() from R Package WGCNA (60), to determine which signature matrix elicited the most meaningful known physiological results.

##### Macrophage subtype marker gene inference

Marker genes for LAM (n=317) and resident macrophages (n=2724) were acquired by using FindMarker() function in *Seurat* (logFC>0.5, adjusted p<0.05) from sub-clustered macrophages (13). Significantly correlated marker genes of LAM or resident macrophages with estimated proportion of macrophages from each publicly available dataset were acquired by using bicorAndPvalue() from R Package WGCNA (60) (p<0.05).

### HUMAN STUDIES AND DECONVOLUTION ANALYSIS

#### Leipzig Childhood Adipose Tissue (LCAT) cohorts

The LCAT cohort includes white children, both female and male, aged 0-18 years who underwent elective orthopedic surgeries, herniotomy/orchidopexy, or other surgical procedures (35). Youth participants with severe diseases and medications that could influence adipose tissue biology, such as diabetes, generalized inflammation, malignant diseases, genetic syndromes, or permanent immobilization, were excluded. Obesity was defined by cutoffs of 1.88 standard deviation score, corresponding to the 97th percentile. During surgery, subcutaneous adipose tissue samples were collected, washed three times in PBS, and immediately frozen in liquid nitrogen for RNA extraction. RNA sequencing was completed on 35 normal weight participants (13 female and 22 male) and 26 participants with obesity (14 female and 12 male) as described previously (61). Gene count matrix was acquired from GSE141221, and subsequently normalized using DESeq2.

#### Metabolically healthy lean (MHL), healthy obese (MHO), and unhealthy obese (MUO) cohorts

The study cohort includes 55 females and males, aged 18-55 years who were classified into three groups based on cardiometabolic health criteria (20). MHL (n = 15; 7 males and 8 females) was defined as having a body mass index (BMI) of 18.5–24.9 kg/m^2^, and normal fasting plasma glucose (<100 mg/dL), oral glucose tolerance (2-h glucose <140 mg/dL), IHTG content (≤5%), plasma triglycerides (<150 mg/dL), and normal whole-body insulin sensitivity, defined as the glucose infusion rate (GIR) per kg fat-free mass divided by the plasma insulin concentration (GIR/I) during the final 20 min of the HECP >40 (μg/kg FFM/min)/(μU/mL). MHO (n = 20; 3 males and 17 females) was defined as having a BMI of 30.0–49.9 kg/m^2^ and normal fasting plasma glucose, oral glucose tolerance, plasma triglycerides, and IHTG content and normal whole-body insulin sensitivity. MUO (n = 20; 3 males and 17 females) was defined as having a BMI of 30.0–49.9 kg/m^2^, impaired fasting glucose or oral glucose tolerance, high IHTG content (≥5%) and impaired whole-body insulin sensitivity, defined as a GIR/I < 40 (μg/kg FFM/min)/(μU/mL). There were no differences in BMI, body fat%, fat-free mass, or subcutaneous abdominal adipose tissue volume between the MHO and MUO, and these two groups were matched for sex and age. Abdominal subcutaneous adipose tissue was collected from the periumbilical area by aspiration using a 3-mm liposuction cannula (Tulip Medical Products, San Diego, CA) connected to a 60cc syringe. Samples were immediately rinsed in ice-cold saline, flash frozen in liquid nitrogen. RNA extraction and sequencing were completed on 15 MHL, 19 MHO, and 19 MUO as described previously (20). Gene count matrix was acquired from GSE244118 and subsequently normalized using DEseq2.

#### MD lipolysis cohorts

The study cohort includes 27 white females (aged 54-70 years) and males (aged 50-70 years), who were classified into three groups (21). Lean (n=9; 6 females and 3 males) was defined as having a BMI of 18-25 kg/m^2^, normal fasting glucose (fP-Glucose < 6.1mmol/l), HbA1c (<42 mmol/mol), and fasting insulin (fS-Insulin < 7.0mU/l). Individuals with obesity with insulin resistance (Obese IR) (n=9; 5 females and 4 males) was defined as having a BMI of 30-40 kg/m^2^, fP-Glucose < 7.0 mmol/l, HbA1c (<48mmol/mol), and fS-Insulin (≥9.0 mU/l). Individuals with obesity with type 2 diabetes (Obese T2D) (n=9; 5 females and 4 males) had BMI of 30-40 kg/m^2^, and have been diagnosed with T2D less than 6 years. Obese NGT and Obese T2D were matched for age, sex, menopausal status, BMI, and fat mass. Abdominal subcutaneous adipose tissue was collected from the periumbilical area by needle aspiration. RNA extraction and sequencing were completed on 9 Lean, 8 MHO (4 females and 4 males), and 8 MUO (4 females and 4 males) as described previously (21). Gene count matrix was acquired from GSE141432 and subsequently normalized using DEseq2.

#### Comprehensive Assessment of Long-term Effects of Reducing Intake of Energy (CALERIE) cohorts

CALERIE cohort includes healthy men (aged 21-50 years) and premenopausal women (aged 21-47 years) without obesity (BMI, 22-27.9 kg/m^2^) who were enrolled in a randomized, controlled trial that targeted to evaluate the time-course effects of 25% calorie restriction (CR) below the subject’s baseline level over a 24 months period. Recruited participants were randomized into either an ad libitum (AL) control group or CR group. RNA-Seq was completed on 13 AL participants and 23 CR participants (62). In the AL group, RNAseq was conducted on 11 participants at month 12 and 6 participants at month 24. In the CR group, RNA-Seq was conducted on 23 participants at month 12 and 12 participants at month 24. Detailed subject characteristics are provided in Table S1.

#### Diet, Obesity, and Genes (DiOGenes) study cohorts

DiOGenes cohort includes adults with overweight/obesity as having a BMI of 27-45 kg/m^2^ and aged 18-65 years (63). 220 participants underwent low-calorie-diet (LCD) period for 8 weeks. During the 8-week weight-loss phase, participants received an LCD that provided 3.3 MJ (800 kcal) per day with the use of Modifast products (Nutrition et Santé). Participants were allowed to eat up to 400 g of vegetables, providing a total, including the LCD, of 3.3 to 4.2 MJ (800 to 1000 kcal) per day. Abdominal subcutaneous adipose tissue biopsies were obtained by needle aspiration, about 10 cm from the umbilicus, under local anesthesia after an overnight fast. Samples were obtained at baseline and upon weight loss. RNA extraction and sequencing were completed on samples as described previously (26). Gene count matrix was acquired from GSE1412221 and subsequently normalized using DEseq2.

#### Very-low-calorie diet (VLCD) cohorts

10 postmenopausal females with obesity (age: 61±4 years) having a BMI >35 kg/m^2^ underwent VLCD that induced approximately 10% of weight loss (27). The VLCD consisted of a commercially available diet (New Direction Program, Robard Corp., Mount Laurel, NJ) that provided ∼800 kcal per day with an estimated macronutrient energy distribution of 54% protein, 26% carbohydrate, 20% fat (including 4% saturated fat and 200 mg of cholesterol), and 10g of fiber. Baseline abdominal subcutaneous adipose tissue biopsy specimen was taken in the left lower quadrant of the abdomen of each subject, whereas the post weight-loss biopsy specimen was taken in the right lower quadrant abdomen. RNA extraction and sequencing were completed as described previously (27). DESeq2-normalized gene count matrix was acquired from GSE106289.

### STATISTICS

Pearson’s correlation coefficient was used to calculate all correlation analyses. Two-tailed independent Student’s t-test was used to compare LCAT cohorts with obesity vs. lean. One-way ANOVA was used to compare MHL vs. MHO vs. MUO and Lean vs. Obese-IR vs. Obese-T2D. Two-way ANOVA linear mixed model was applied to examine the main effects of time, group, and time x group interaction effects from CALERIE cohorts (time, Baseline vs. Year1 vs. Year2; group, AL vs. CR). For significant ANOVA results, post hoc pairwise comparisons were performed using the estimated marginal means with Tukey’s adjustment for multiple comparisons. Two-tailed paired Student’s T test was used to examine the effect of DIWL in DiOGenes cohorts and women with obesity in Aleman et al., 2017. Statistical computations were performed using R (R, Vienna, Austria). P value < 0.05 was considered statistically significant.

## DATA AND CODE AVAILABILITY

Gene signature matrix and top 6000 HVG list have been uploaded to https://github.com/KWhytock13/deconvolution-wat.

Code for generating source data and running the deconvolution pipeline is also available at https://github.com/KWhytock13/deconvolution-wat.

## ACKNOWLEDGMENTS

This study was supported by The National Institutes of Health (R01 AG066474, U01 AR071133 supplement). Data from the CALERIE analysis was supported by the National Institutes of Health **(R01 AG061378, R33 AG070455)**

## AUTHOR CONTRIBUTIONS

CA, MMS, LMS, and KW designed the study. All authors contributed to data acquisition, analysis, and interpretation. CA, LMS, and KW drafted the work. All authors have participated in revising the work and approved the final version of the manuscript. We would like to thank Dr. Daniel Belsky, Dr Kim Huffman and Dr Calen Ryan for sharing CALERIE RNA-Seq data for our analyses.

---

**Table S1.** CALERIE^TM^ subject characteristics. Basic subject characteristics from CALERIE^TM^ participants whose ASAT samples were sequenced and used for deconvolution. a, significantly different against AL-Baseline; b, significantly different against AL-Month 12; c, significantly different against AL-Month 24. d, significantly different against CR-Baseline. AL, Ad Libitum; CR, Calorie Restriction, BMI, Body Mass Index.

|  | AL |  |  | CR |  |  |
| --- | --- | --- | --- | --- | --- | --- |
| Time point | Baseline<br>(n=14) | Month 12<br>(n=11) | Month 24<br>(n=6) | Baseline<br>(n=23) | Month 12<br>(n=23) | Month 24<br>(n=12) |
| Age (y) | 38 ± 8 | NA | NA | 37 ± 7 | NA | NA |
| Sex | 8F, 6M | 5F, 6M | 3F, 3M | 18F, 5M | 18F, 5M | 9F, 3M |
| BMI (kg/m<sup>2</sup>) | 25.1 ± 1.4 | 25.1 ± 1.9 | 26.0 ± 2.4 | 25.5 ± 1.6 | 22.4 ± 1.6<sup>a,b,c,d</sup> | 22.6 ± 1.9<sup>a,b,c,d</sup> |
| Weight (kg) | 74.6 ± 9.7 | 75.3 ± 11.6 | 78.0 ± 8.0 | 71.5 ± 9.2 | 63.0 ± 8.9<sup>a,b,c,d</sup> | 63.5 ± 9.1<sup>a,b,c,d</sup> |
| Δ Fat mass (kg) | NA | 0.3 ± 1.7 | 0.2 ± 1.9 | NA | -6.2 ± 2.0<sup>b,c</sup> | -6.5 ± 1.9<sup>b,c</sup> |
| Δ Fat-free mass (kg) | NA | 0.1 ± 0.9 | 0.9 ± 1.3 | NA | -2.0 ± 1.2<sup>b,c</sup> | -1.8 ± 1.1<sup>b,x</sup> |

**Table S2.** Subject characteristics of DioGenes and Aleman et al., 2017.

|  | DiOgenes<br>(8-week LCD) | Aleman et al., 2017<br>(10% Weight loss targeted VLCD) |
| --- | --- | --- |
| Age (y) | 41 ± 6 | 61 ± 4 |
| Baseline BMI (kg/m<sup>2</sup>) | 34.8 ± 4.9 | 38.8 ± 3.4 |
| Weight loss during intervention (%) | -11.1 ± 2.7 | -10.3 |
| Intervention Duration (weeks) | ~8 | 6.6 ± 2.2 |

**Figure S1.**
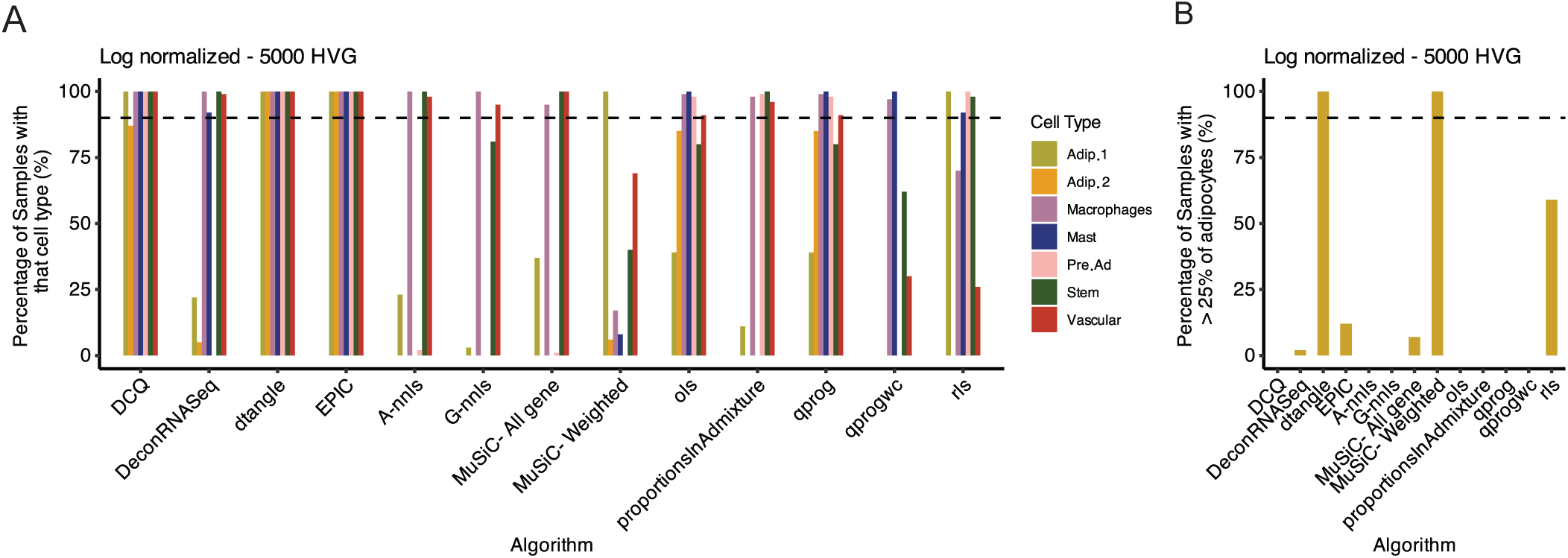
The capability of different algorithms to predict cell types in bulk RNA-Seq data from METSIM and normalized counts from snRNA-Seq. (A) The percentage of samples from the METSIM cohort that are estimated to contain that cell type when predicted with different deconvolution algorithms using log normalized signature matrix subset to 5000 HVG. (B) The percentage of samples from the METSIM cohort that are estimated to have >25% of adipocytes predicted with different deconvolution algorithms using log normalized signature matrix subset to 5000 HVG.

**Figure S2.**
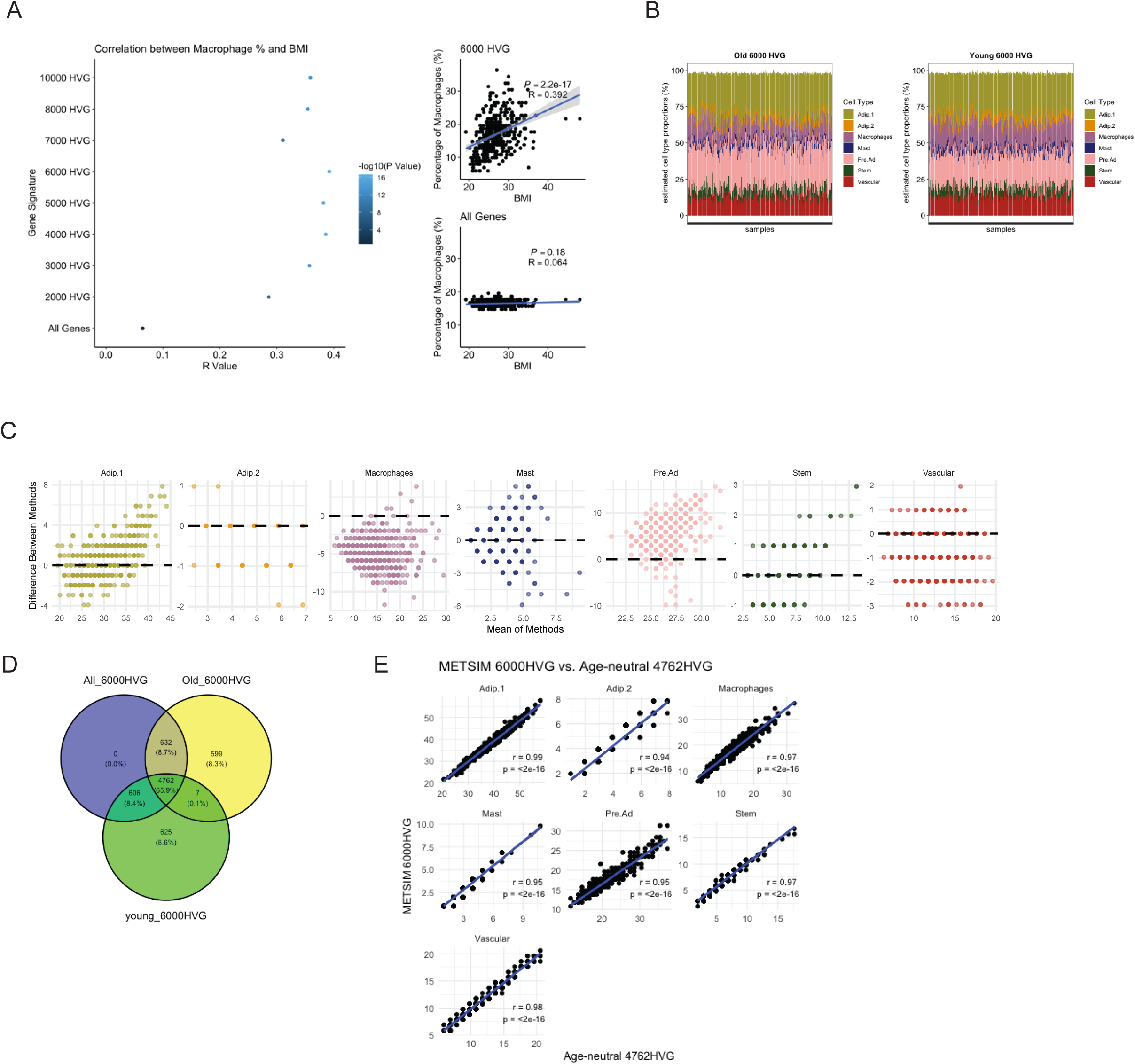
HVG optimization. (A) Estimated cell type proportions of each METSIM bulk RNA-Seq samples using different HVG to modify the signature matrix. Algorithm dtangle was used and the data was log normalized. R value and –log10(p value) for estimated macrophage proportion and BMI for the METSIM bulk RNA-Seq data. Scatterplot showing the estimated macrophage proportion and BMI for the 6000 HVG or all genes signature. (B) Deconvolution result of METSIM bulk RNA-seq using age-group-specific gene signature matrice and 6000 HVG. (C) Bland-Altman plots for estimated proportion of each cell type using Older (n=10) vs. young (n=10) snRNAseq data. Contrast is Older - Younger. Points above dashed line (Y=0) indicates higher estimation of cell proportion when using reference data from Older individuals. (D) Venn diagram of overlapping HVG among integrated 6000 HVG, Old-specific 6000 HVG, and Young-specific 6000 HVG. (E) Scatter plot for each cell type showing correlation between estimated cell proportion in METSIM using integrated 6000 HVG and ‘age-neutral’ 4762 HVG.

## References

1. Kershaw EE, and Flier JS. Adipose tissue as an endocrine organ. The Journal of Clinical Endocrinology & Metabolism. 2004;89(6):2548–56.

2. Goodpaster BH, and Sparks LM. Metabolic Flexibility in Health and Disease. Cell Metab. 2017;25(5):1027–36.

3. Trayhurn P. Endocrine and signalling role of adipose tissue: new perspectives on fat. Acta Physiologica Scandinavica. 2005;184(4):285–93.

4. Eto H, Suga H, Matsumoto D, Inoue K, Aoi N, Kato H, et al. Characterization of structure and cellular components of aspirated and excised adipose tissue. Plastic and reconstructive surgery. 2009;124(4):1087–97.

5. Corvera S. Cellular Heterogeneity in Adipose Tissues. Annu Rev Physiol. 2021;83:257–78.

6. Frayn KN. Adipose tissue as a buffer for daily lipid flux. Diabetologia. 2002;45(9):1201–10.

7. Goossens GH. The Metabolic Phenotype in Obesity: Fat Mass, Body Fat Distribution, and Adipose Tissue Function. Obes Facts. 2017;10(3):207–15.

8. Klöting N, Fasshauer M, Dietrich A, Kovacs P, Schön MR, Kern M, et al. Insulin-sensitive obesity. American Journal of Physiology-Endocrinology and Metabolism. 2010;299(3):E506–E15.

9. Brotman SM, Oravilahti A, Rosen JD, Alvarez M, Heinonen S, van der Kolk BW, et al. Cell-Type Composition Affects Adipose Gene Expression Associations With Cardiometabolic Traits. Diabetes. 2023;72(11):1707–18.

10. Schleh MW, Ameka M, Rodriguez A, and Hasty AH. Deficiency of the hemoglobin-haptoglobin receptor, CD163, worsens insulin sensitivity in obese male mice. bioRxiv. 2024.

11. Kosteli A, Sugaru E, Haemmerle G, Martin JF, Lei J, Zechner R, and Ferrante AW, Jr. Weight loss and lipolysis promote a dynamic immune response in murine adipose tissue. J Clin Invest. 2010;120(10):3466–79.

12. Aron-Wisnewsky J, Tordjman J, Poitou C, Darakhshan F, Hugol D, Basdevant A, et al. Human adipose tissue macrophages: m1 and m2 cell surface markers in subcutaneous and omental depots and after weight loss. The Journal of Clinical Endocrinology & Metabolism. 2009;94(11):4619–23.

13. Whytock KL, Divoux A, Sun Y, Pino MF, Yu G, Jin CA, et al. Aging human abdominal subcutaneous white adipose tissue at single cell resolution. Aging Cell. 2024:e14287.

14. Emont MP, Jacobs C, Essene AL, Pant D, Tenen D, Colleluori G, et al. A single-cell atlas of human and mouse white adipose tissue. Nature. 2022;603(7903):926-33.

15. Massier L, Jalkanen J, Elmastas M, Zhong J, Wang T, Nono Nankam PA, et al. An integrated single cell and spatial transcriptomic map of human white adipose tissue. Nat Commun. 2023;14(1):1438.

16. Raulerson CK, Ko A, Kidd JC, Currin KW, Brotman SM, Cannon ME, et al. Adipose Tissue Gene Expression Associations Reveal Hundreds of Candidate Genes for Cardiometabolic Traits. Am J Hum Genet. 2019;105(4):773–87.

17. Lee M-J, Wu Y, and Fried SK. Adipose tissue heterogeneity: implication of depot differences in adipose tissue for obesity complications. Molecular aspects of medicine. 2013;34(1):1–11.

18. Weisberg SP, McCann D, Desai M, Rosenbaum M, Leibel RL, and Ferrante AW. Obesity is associated with macrophage accumulation in adipose tissue. The Journal of clinical investigation. 2003;112(12):1796–808.

19. Yang CH, Fagnocchi L, Apostle S, Wegert V, Casani-Galdon S, Landgraf K, et al. Independent phenotypic plasticity axes define distinct obesity sub-types. Nat Metab. 2022;4(9):1150–65.

20. Petersen MC, Smith GI, Palacios HH, Farabi SS, Yoshino M, Yoshino J, et al. Cardiometabolic characteristics of people with metabolically healthy and unhealthy obesity. Cell Metab. 2024;36(4):745–61 e5.

21. Fryk E, Olausson J, Mossberg K, Strindberg L, Schmelz M, Brogren H, et al. Hyperinsulinemia and insulin resistance in the obese may develop as part of a homeostatic response to elevated free fatty acids: A mechanistic case-control and a population-based cohort study. EBioMedicine. 2021;65:103264.

22. Crewe C, An YA, and Scherer PE. The ominous triad of adipose tissue dysfunction: inflammation, fibrosis, and impaired angiogenesis. The Journal of clinical investigation. 2017;127(1):74–82.

23. Pasarica M, Sereda OR, Redman LM, Albarado DC, Hymel DT, Roan LE, et al. Reduced adipose tissue oxygenation in human obesity: evidence for rarefaction, macrophage chemotaxis, and inflammation without an angiogenic response. Diabetes. 2009;58(3):718–25.

24. Bluher M. Adipose tissue dysfunction contributes to obesity related metabolic diseases. Best Pract Res Clin Endocrinol Metab. 2013;27(2):163–77.

25. Magkos F, Fraterrigo G, Yoshino J, Luecking C, Kirbach K, Kelly SC, et al. Effects of Moderate and Subsequent Progressive Weight Loss on Metabolic Function and Adipose Tissue Biology in Humans with Obesity. Cell Metab. 2016;23(4):591–601.

26. Imbert A, Vialaneix N, Marquis J, Vion J, Charpagne A, Metairon S, et al. Network Analyses Reveal Negative Link Between Changes in Adipose Tissue GDF15 and BMI During Dietary-induced Weight Loss. J Clin Endocrinol Metab. 2022;107(1):e130–e42.

27. Aleman JO, Iyengar NM, Walker JM, Milne GL, Da Rosa JC, Liang Y, et al. Effects of Rapid Weight Loss on Systemic and Adipose Tissue Inflammation and Metabolism in Obese Postmenopausal Women. J Endocr Soc. 2017;1(6):625–37.

28. Fritzen AM, Lundsgaard AM, Jordy AB, Poulsen SK, Stender S, Pilegaard H, et al. New Nordic Diet-Induced Weight Loss Is Accompanied by Changes in Metabolism and AMPK Signaling in Adipose Tissue. J Clin Endocrinol Metab. 2015;100(9):3509–19.

29. Rickman AD, Williamson DA, Martin CK, Gilhooly CH, Stein RI, Bales CW, et al. The CALERIE Study: design and methods of an innovative 25% caloric restriction intervention. Contemp Clin Trials. 2011;32(6):874–81.

30. Russo L, and Lumeng CN. Properties and functions of adipose tissue macrophages in obesity. Immunology. 2018;155(4):407–17.

31. Åkra S, Aksnes TA, Flaa A, Eggesbø HB, Opstad TB, Njerve IU, and Seljeflot I. Markers of remodeling in subcutaneous adipose tissue are strongly associated with overweight and insulin sensitivity in healthy non-obese men. Scientific Reports. 2020;10(1):14055.

32. Mozaffarian D, Hao T, Rimm EB, Willett WC, and Hu FB. Changes in diet and lifestyle and long-term weight gain in women and men. New England journal of medicine. 2011;364(25):2392–404.

33. Tam CS, Tordjman J, Divoux A, Baur LA, and Clement K. Adipose tissue remodeling in children: the link between collagen deposition and age-related adipocyte growth. J Clin Endocrinol Metab. 2012;97(4):1320–7.

34. Spalding KL, Arner E, Westermark PO, Bernard S, Buchholz BA, Bergmann O, et al. Dynamics of fat cell turnover in humans. Nature. 2008;453(7196):783–7.

35. Landgraf K, Rockstroh D, Wagner IV, Weise S, Tauscher R, Schwartze JT, et al. Evidence of early alterations in adipose tissue biology and function and its association with obesity-related inflammation and insulin resistance in children. Diabetes. 2015;64(4):1249–61.

36. Ravussin E, and Smith SR. Increased fat intake, impaired fat oxidation, and failure of fat cell proliferation result in ectopic fat storage, insulin resistance, and type 2 diabetes mellitus. Ann N Y Acad Sci. 2002;967:363–78.

37. Cypess AM. Reassessing human adipose tissue. New England Journal of Medicine. 2022;386(8):768–79.

38. Shulman GI. Ectopic fat in insulin resistance, dyslipidemia, and cardiometabolic disease. N Engl J Med. 2014;371(12):1131–41.

39. Nicklas BJ, Ambrosius W, Messier SP, Miller GD, Penninx BW, Loeser RF, et al. Diet-induced weight loss, exercise, and chronic inflammation in older, obese adults: a randomized controlled clinical trial. Am J Clin Nutr. 2004;79(4):544–51.

40. Christiansen T, Paulsen SK, Bruun JM, Pedersen SB, and Richelsen B. Exercise training versus diet-induced weight-loss on metabolic risk factors and inflammatory markers in obese subjects: a 12-week randomized intervention study. Am J Physiol Endocrinol Metab. 2010;298(4):E824–31.

41. Meydani SN, Das SK, Pieper CF, Lewis MR, Klein S, Dixit VD, et al. Long-term moderate calorie restriction inhibits inflammation without impairing cell-mediated immunity: a randomized controlled trial in non-obese humans. Aging (Albany NY). 2016;8(7):1416.

42. Capel F, Klimcakova E, Viguerie N, Roussel B, Vitkova M, Kovacikova M, et al. Macrophages and adipocytes in human obesity: adipose tissue gene expression and insulin sensitivity during calorie restriction and weight stabilization. Diabetes. 2009;58(7):1558–67.

43. Asterholm IW, Tao C, Morley TS, Wang QA, Delgado-Lopez F, Wang ZV, and Scherer PE. Adipocyte inflammation is essential for healthy adipose tissue expansion and remodeling. Cell metabolism. 2014;20(1):103–18.

44. Murphy J, Moullec G, and Santosa S. Factors associated with adipocyte size reduction after weight loss interventions for overweight and obesity: a systematic review and meta-regression. Metabolism. 2017;67:31–40.

45. Ejaz A, Mitterberger MC, Lu Z, Mattesich M, Zwierzina ME, Horl S, et al. Weight Loss Upregulates the Small GTPase DIRAS3 in Human White Adipose Progenitor Cells, Which Negatively Regulates Adipogenesis and Activates Autophagy via Akt-mTOR Inhibition. EBioMedicine. 2016;6:149–61.

46. Rossmeislova L, Malisova L, Kracmerova J, Tencerova M, Kovacova Z, Koc M, et al. Weight loss improves the adipogenic capacity of human preadipocytes and modulates their secretory profile. Diabetes. 2013;62(6):1990–5.

47. Lofgren P, Andersson I, Adolfsson B, Leijonhufvud BM, Hertel K, Hoffstedt J, and Arner P. Long-term prospective and controlled studies demonstrate adipose tissue hypercellularity and relative leptin deficiency in the postobese state. J Clin Endocrinol Metab. 2005;90(11):6207–13.

48. Prins JB, Walker NI, Winterford CM, and Cameron DP. Human adipocyte apoptosis occurs in malignancy. Biochemical and biophysical research communications. 1994;205(1):625–30.

49. Fischer-Posovszky P, Wang QA, Asterholm IW, Rutkowski JM, and Scherer PE. Targeted deletion of adipocytes by apoptosis leads to adipose tissue recruitment of alternatively activated M2 macrophages. Endocrinology. 2011;152(8):3074–81.

50. Sabina Pfister VK, Enrico Ferrero. granulator: Rapid benchmarking of methods for *in silico* deconvolution of bulk RNA-seq data. 2024.

51. Monaco G, Lee B, Xu W, Mustafah S, Hwang YY, Carre C, et al. RNA-Seq Signatures Normalized by mRNA Abundance Allow Absolute Deconvolution of Human Immune Cell Types. Cell Rep. 2019;26(6):1627–40 e7.

52. Gong T, and Szustakowski JD. DeconRNASeq: a statistical framework for deconvolution of heterogeneous tissue samples based on mRNA-Seq data. Bioinformatics. 2013;29(8):1083–5.

53. Abbas AR, Wolslegel K, Seshasayee D, Modrusan Z, and Clark HF. Deconvolution of blood microarray data identifies cellular activation patterns in systemic lupus erythematosus. PLoS One. 2009;4(7):e6098.

54. Hunt GJ, Freytag S, Bahlo M, and Gagnon-Bartsch JA. dtangle: accurate and robust cell type deconvolution. Bioinformatics. 2019;35(12):2093–9.

55. Danziger SA, Gibbs DL, Shmulevich I, McConnell M, Trotter MWB, Schmitz F, et al. ADAPTS: Automated deconvolution augmentation of profiles for tissue specific cells. PLoS One. 2019;14(11):e0224693.

56. Altboum Z, Steuerman Y, David E, Barnett-Itzhaki Z, Valadarsky L, Keren-Shaul H, et al. Digital cell quantification identifies global immune cell dynamics during influenza infection. Mol Syst Biol. 2014;10(2):720.

57. Wang X, Park J, Susztak K, Zhang NR, and Li M. Bulk tissue cell type deconvolution with multi-subject single-cell expression reference. Nat Commun. 2019;10(1):380.

58. Racle J, and Gfeller D. EPIC: a tool to estimate the proportions of different cell types from bulk gene expression data. Bioinformatics for cancer immunotherapy: methods and protocols. 2020:233–48.

59. Laakso M, Kuusisto J, Stancakova A, Kuulasmaa T, Pajukanta P, Lusis AJ, et al. The Metabolic Syndrome in Men study: a resource for studies of metabolic and cardiovascular diseases. J Lipid Res. 2017;58(3):481–93.

60. Langfelder P, and Horvath S. WGCNA: an R package for weighted correlation network analysis. BMC Bioinformatics. 2008;9:559.

61. Dalgaard K, Landgraf K, Heyne S, Lempradl A, Longinotto J, Gossens K, et al. Trim28 Haploinsufficiency Triggers Bi-stable Epigenetic Obesity. Cell. 2016;164(3):353–64.

62. Ryan CP, Corcoran DL, Banskota N, Eckstein IC, Floratos A, Friedman R, et al. The CALERIE() Genomic Data Resource. bioRxiv. 2024.

63. Larsen TM, Dalskov S-M, van Baak M, Jebb SA, Papadaki A, Pfeiffer AF, et al. Diets with high or low protein content and glycemic index for weight-loss maintenance. New England Journal of Medicine. 2010;363(22):2102–13.

